# Stem photosynthesis from wild Prunus arabica enhances growth, advances bloom and increases yield in cultivated almond

**DOI:** 10.64898/2026.06.23.734067

**Authors:** Dan Zeira, Ori Eisenbach, Rotem Harel-Beja, Taly Trainin, Kamel Hatib, Lotem Terner, Mohamed Abd-Elhadi, Hillel Brukental, Or Shapira, Yotam Zait, Doron Holland, Tamar Azoulay-Shemer

## Abstract

Rising winter temperatures threaten deciduous fruit tree productivity by depleting carbohydrate reserves during dormancy. This study investigated Stem Photosynthetic Capacity (SPC), a rare adaptive trait from wild *Prunus arabica*, as a mechanism to enhance almond carbon economy. Using extreme segregating groups from the F1 population (*P. dulcis* X *P. arabica*), we evaluated physiological performance through high-resolution lysimetric and multi-year orchard monitoring. High-SPC [SPC(+)] genotypes maintained significantly greater stem CO_2_ assimilation and transpiration during leafless periods compared to low-SPC [SPC(-)] progenies. Over five successive seasons, SPC(+) trees exhibited a 33.3% increase in trunk secondary growth and reached 10% bloom approximately 8 days earlier. Most importantly, the SPC(+) group achieved a 4.6-fold increase in mean kernel yield when compared to SPC(-) group. These findings demonstrate that SPC provides a flexible, supplementary winter carbon source that directly supports both vegetative and reproductive development. Integrating SPC into commercial almond breeding programs may offer a valuable strategy to improve climate resilience and help sustain yields under warming conditions.

**Highlight:** Integrating stem photosynthesis into commercial almond hybrids provides a winter carbon source that advances blooming, expands trunk growth by ∼33%, and increases kernel yields by more than 4.5-fold.

## 1. Introduction

Almonds (*Prunus dulcis* (Mill.) D. A. Webb) are among the most globally significant crops, recognized for their high nutritional and health benefits (Estruch et al., 2018; Hyson et al., 2002; King et al., 2008; Martinez-Gonzalez et al., 2008). Deciduous fruit trees, such as almonds, enter dormancy at the onset of winter and resume growth only after exposure to a species-specific period of low temperatures, known as chilling requirements (CR). Adequate chilling is essential for proper bud differentiation and synchronization of early spring flowering, which ensures efficient pollination, fruit set, and development (Sanchez-Perez et al., 2014). A primary concern regarding global climate change is the impact of rising winter temperatures (Luedeling et al., 2011). Elevated winter temperatures in deciduous fruit trees can modify dormancy progression and carbohydrate metabolism in dormant organs, increasing metabolic activity and respiratory demand and thereby enhancing the consumption of stored carbohydrates (Sperling et al., 2019). This temperature-driven shift in starch synthesis and degradation can limit the accumulation of soluble carbohydrates in flower buds, so that carbohydrate availability becomes insufficient during the critical phases of flowering and fruit set (Fadon et al., 2018; Fernandez et al., 2018). Such physiological disruptions have been associated with abnormal bloom behavior, impaired flower development, which eventually leads to reduced fruit set, and lower yields (Tominaga et al., 2021).

These climatic trends highlight the urgent need to enhance the physiological resilience of deciduous crops, enabling trees to better adapt to changing environmental conditions. Throughout agricultural development and breeding programs, many valuable traits preserved in wild almond varieties have been lost from commercial cultivars. In a preliminary study, we explored the rare wild almond species, “Arabica almond” (*Amygdalus arabica Olivier, P. arabica*), which grows in desert regions surrounding the Fertile Crescent. This species thrives under extreme conditions, characterized by drought and significant temperature fluctuations. *P. arabica* exhibits a shrub-like architecture rather than a tree, possesses an extensive root system, and is widely recognized as drought tolerance (El-Sheikh et al., 2019; Sorkheh et al., 2011).

We identified a unique gas exchange and carbon assimilation strategy in this species that distinctly differs from the characteristics of commercial almond varieties (Brukental et al., 2021; Trainin et al., 2022). Our preliminary findings demonstrate that the stems of *P. arabica*, which account for 60% of the tree’s surface area, remain green year-round, including during the winter when the tree is otherwise leafless. Further investigation revealed that these stems perform gas exchange, similar to leaves, through functional stomata that regulate transpiration and photosynthetic carbon dioxide assimilation in response to environmental changes(Trainin et al., 2022).

The ability to photosynthesize through the stem is characteristic of desert plants adapted to extreme conditions; these plants may shed their leaves to reduce water loss yet continue to assimilate CO_2_ through the green stems (Avila-Lovera et al., 2019). This unique trait, known as Stem Photosynthetic Capacity (SPC), provides an additional carbon source for the tree and contributes to the non-structural carbohydrate pool, with implications for xylem hydraulic safety (Natale et al., 2023). To elucidate the role of the SPC trait in almond tree development, an F1 experimental population was established in the almond orchard at the Newe Ya’ar Research Center. This population was derived from a cross between the commercial almond cultivar ‘Um-El-Fahem’ (*P. dulcis*) and the wild almond species *P. arabica* (UEF X *P. arabica*), and consists of approximately 90 progenies (Brukental et al., 2021). The SPC trait segregates within the population, with each progeny displaying a different level of SPC expression. The physiological value of the SPC trait lies in providing greater flexibility, enabling deciduous fruit trees, such as almonds, to adapt to changing climate conditions by maintaining a favorable energy balance, consistent with evidence that stem photosynthesis can sustain carbohydrate reserves and modulate xylem vulnerability to embolism (Natale et al., 2023). This is especially important during periods when respiration rates remain high, but energy production by leaves is constrained, or leaves are absent (Avila-Lovera et al., 2019; Nilsen and Bao, 1990), such as during warm, leafless winters, when green stems can continue to assimilate CO_2_.

Based on preliminary data, we propose the following hypotheses: (1) The stem photosynthetic capacity (SPC) observed in the wild species *P. arabica* contributes significantly to the annual carbon gain in cultivated almond trees. (2) The SPC trait is active throughout the year, but its magnitude is modulated by the photosynthetic activity of the primary leaf tissue, which declines during the leafless winter period. (3) This additional carbon source supports tree growth and reproduction, potentially enhancing overall yield.

## 2. Materials and Methods

### 2.1 Plant Material

An F1 population was obtained from a successful cross between two almond species. *P. arabica* (♂) was used to pollinate the commercial variety ‘Um-El-Fahem’ (♀), yielding 95 progenies. We vegetatively reproduced *P. arabica*, ‘Um-El-Fahem’, and the F1 progenies by grafting scions onto Hansen 563 rootstocks. All trees are cultivated in the almond orchard in Newe Ya’ar Research Center in the Yizra’el Valley (latitude 34◦ 420N, longitude 35◦ 110E, Mediterranean temperate to subtropical climate), with one copy for each, and planted in winter 2018.

### 2.2 Plant Experimental System

Parental lines (’Um-El-Fahem’ and *P. arabica*): The parents of the F1 population, the Israeli leading commercial cv. ‘Um-El-Fahem’ (UEF) and the wild almond species, *P. arabica,* were grown in two copies for each, grafted on Hansen rootstock, and planted in winter 2018.

#### F1 Segregating population

The experimental population consists of 12 selected lines from the F1 ‘Um-El-Fahem’ x *P. arabica* population, 6 progenies with the SPC trait and 6 progenies without the parental lines, ‘Um-El-Fahem’ (♀), *P. arabica* (♂). The decision regarding which progeny to select is based on preliminary work (Brukental et al., 2021) in which we characterized, both physiologically and genetically, the segregation of the SPC trait within this F1 population. Throughout all parts of the experiment, tests and measurements were performed on all 12 progenies and the two parental lines.

### Lysimetric Experiment

### 2.3 Plant Material

Multiple replicates (n=3-7) of the selected progenies and their parents were grafted onto Hansen 563 rootstocks in 4-liter pots. The trees were grown at the Newe Ya’ar Research Center in 4-liter pots under standard irrigation and fertilization conditions commonly used in the transplant industry for 6 months. Approximately one month before transferring the plants to the lysimeter system, they were moved to 7-liter pots.

### 2.4 Lysimetric system (ICORE phenotyping platform).

Plants were grown and analyzed during the late winter and the spring seasons (January 2024 to May 2024) in the I CORE lysimetric system, located at the Faculty of Agriculture, Rehovot, Israel. The ICOR lysimetric system is a phenotyping platform for soil-plant-atmosphere continuum under dynamic environmental conditions, composed of highly sensitive, temperature–compensated load cells equipped with gravimetric lysimeters, soil moisture and atmospheric sensors (Halperin et al., 2017). The system measures five quantitative physiological traits (QPTs) simultaneously: (1) whole-plant transpiration rate; (2) daily/periodic increase in plant biomass; (3) canopy stomatal conductance; (4) whole-plant water use efficiency (WUEwp); and (5) soil water content (VWC). Data were recorded every minute throughout the experiment. The plants were grown and analyzed under natural sunlight, with regulated temperatures that closely mirror outdoor conditions (Paul et al., 2023). Throughout the experiment, the plants were grown under well-fertilized and irrigation conditions. The plants were placed in an I-CORE greenhouse in a randomized block design with 3- 7 replicates per progeny. The arrangement of the blocks was managed using the control software of the I-CORE system.

### 2.5 Gas exchange measurements

Gas exchange measurements for both stems and leaves were conducted in the I-CORE greenhouse utilizing a LI-6800 Portable Photosynthesis System equipped with a 2 cm² chamber. Stem assessments were performed from March to May 2024 on 6-month-old trees, while leaf measurements were carried out in March 2024. For each plant, four replicates (n=4 leaves/stems) were analyzed per session, with all data collection occurring between 09:30 and 13:00. The chamber environment was standardized with a CO_2_ reference of 415 ppm and a photosynthetic photon flux density of 1000 µmol m⁻² s⁻¹ (90% red, 10% blue), while temperature and relative humidity were continuously adjusted to match prevailing ambient condition. Stem measurements were specifically localized approximately 15 cm from the tip, and subsequent results were normalized to the stem surface area, calculated as 1.6 cm * π * stem diameter (cm).

### 2.6 Biomass Accumulation and Partitioning

Total plant biomass was determined based on dry weight and categorized into three distinct tissue types: leaves, stems, and the main trunk. To determine the dry mass, harvested plant material from each individual was isolated in labeled paper bags and dehydrated in a forced-air oven at 60°C for 4 days until a constant weight was achieved. Following the drying period, the samples were removed from the oven and allowed to equilibrate to room temperature for approximately one hour to prevent moisture fluctuations during measurement. The final dry weight for each tissue component was quantified using a digital scale.

### Orchard Experiment

### 2.7 Plant Material

A description of the experimental population can be found in Section 2.13

### 2.8 Phenological Measurements

#### 2.81 Trunk circumference

Trunk circumference was recorded annually over five consecutive years. Measurements were conducted each autumn (November–December) using a flexible measuring tape to encircle the trunk. To ensure consistency across all progenies, the measurement was taken at a standardized point approximately 10 cm above the graft union.

#### 2.82 Phenological Assessments

Bloom phenology was monitored annually over four consecutive years. Observations commenced in late winter and continued until the trees reached full bloom. The specific timing of flowering was defined as the date tree reached 10% bloom intensity. To ensure accuracy, visual inspections of the entire canopy were conducted on a bi-daily basis throughout the flowering period.

#### 2.83 Yield Determination

Annual fruit yield was recorded over five year. Harvesting was performed manually at the end of each growing season, strictly adhering to commercial cultivation protocols. The total yield per tree was quantified as clean crop weight, which comprised both the kernel (seed) and the surrounding hull (husk).

### 2.9 Physiological Measurements Orchard Gas Exchange Measurements

CO_2_ assimilation rates for both stems and leaves were measured in the Newe Ya’ar almond orchard utilizing the same equipment and parameters described for the ICOR greenhouse experiments (see section 2.5). These field measurements were conducted from December 2024 through August 2025 on one-year-old stems. While environmental conditions were consistent with previous protocols, the leaf analysis was adjusted to include three leaves (n = 3) per tree during each measurement session. All measurements were performed on the previously described experimental population to ensure comparative consistency across the study.

## 3. Results

### High-Resolution Physiological Monitoring via Lysimetry

The implementation of a gravimetric lysimetric system facilitated high-resolution, continuous monitoring of whole-tree physiological behavior. This platform enabled the detection of subtle gas exchange and transpiration dynamics under controlled environmental conditions, over the experimental period.

### 3.1 Leaf and Stem CO_2_ Assimilation Dynamics

To evaluate the relative contribution of different tissues to the total carbon budget, net CO_2_ assimilation rates (*A_n_*) were quantified for leaves (*A*_leaf_) and stems (*A*_stem_) across all twelve progenies and their parental lines before and after defoliation. During the springtime (March 2024), measurements were conducted on both leaves (Fig.1) and stems (Fig. 1B), while after leaf removal, only stems were measured (Fig. 1B).

**Fig 1.**
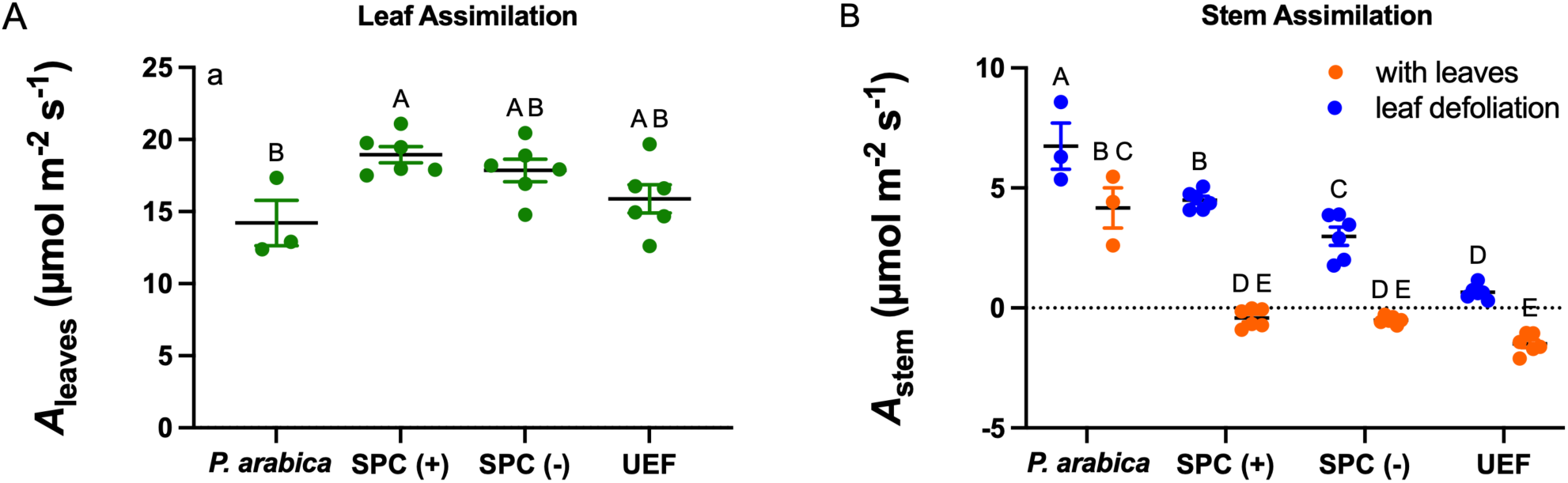
Leaf and Stem CO_2_ Assimilation rate (*A_n_*) of ‘Um-El-Fahem’ x *P. arabica* F1 progenies and their parental lines. Net CO₂ assimilation rates (*A_n_*) of ‘Um-El-Fahem’ x *P. arabica* F1 progenies and their parental lines measured during spring 2024 using a LI-6800 portable photosynthesis system. Experimental groups include F1 progenies with the stem photosynthetic capability trait (SPC(+), n = 6), progenies lacking the trait (SPC(-), n = 6), and the parental lines *P. arabica* (n = 3) and ‘Um-El-Fahem’ (UEF, n = 7). **(A)** *A*_leaf_ measured during spring 2024. Data represent mean values +/- STDEV. Different letters indicate significant differences among groups based on [Nonparametric comparisons for each pair using Wilcoxon method, *P <* 0.05]. **(B)** *A*_stem_ measured in the same F1 progenies and parental lines before (with leaves) and after artificial leaf removal (without leaves). Data represent mean values +/-STDEV. Lowercase letters denote significant differences among groups in the foliated state, whereas uppercase letters denote significant differences in the defoliated state based on [Nonparametric comparisons for each pair using the Wilcoxon method, *P <* 0.05]. Statistically significant differences between foliated and defoliated states within each group are indicated.

During the spring, *A*_leaf_ values were relatively uniform across all F1 progenies, with no significant differences observed between SPC(+) and SPC(-) progenies. Nevertheless, both F1 groups exhibited higher leaf assimilation rates than the parental lines, ‘Um-El-Fahem’ and *P. arabica* (Fig. 1A). Specifically, SPC(+) progenies showed significantly higher *A*_leaf_ compared to both parents, whereas SPC(-) progenies displayed intermediate values that did not differ significantly from either group. In contrast to the relatively homogeneous leaf responses, stem photosynthetic performance exhibited marked differences depending on foliation status. Under foliated spring conditions, *A*_stem_ values in all 12 F1 progenies and in the ‘Um-El-Fahem’ parent were negligible, ranging from 0.4 to -1.4 µmol CO₂ m^-^² s^-^¹, while the wild parent *P. arabica* maintained a significantly positive *A*_stem_ of 4.2 µmol CO₂ m^-^² s^-^¹ (Fig.1B). Following artificial defoliation, *A*_stem_ increased across all F1 progenies, with the strongest response observed in *P. arabica*, where assimilation rates increased 1.5-fold from 4.2 to 6.1 µmol CO₂ m^-^² s^-^¹ (*P* = 0.002). Among the defoliated F1 progenies, SPC(+) individuals exhibited significantly higher stem assimilation rates (mean = 4.5 µmol CO₂ m^-^² s⁻¹) compared to SPC(-) progenies (mean = 3.2 µmol CO₂ m^-^² s^-^¹; *P* = 0.008). The commercial parent ‘Um-El-Fahem’ showed a significant physiological response to leaf fall, shifting to low positive *A*_stem_ (-0.8 μmol CO₂ m^-^¹ s^-^¹) in contrast to its foliage state (0.65 μmol CO₂ m^-^¹ s) (Fig. 1B).

### 3.2 Physiological Regulation of Whole-Tree Transpiration and Conductance

To evaluate the impact of the Stem Photosynthetic Capability (SPC) trait on whole-plant water use and assess active physiological regulation, agronomic, whole-tree daily transpiration was monitored continuously throughout the late winter and spring using the gravimetric lysimetric system. The daily water consumption was quantified in ml of water per day served as the primary measure of whole-canopy daily transpiration rate.

As the trees emerged from winter dormancy during week 7, they initiated bud break and leaf flush, leading to a marked increase in transpiration across all progenies. A significant physiological shift was observed following artificial defoliation in week 11. By week 12, when all trees were in a completely leafless state, transpiration was driven almost exclusively by stem-based water loss. During this period, multiple comparison analysis revealed significant differences in agronomic transpiration rates among the experimental groups (Fig. 2). Significant differences were observed between parental and progeny agronomic transpiration rates during the leafless state. Specifically, the wild parent *P. arabica* exhibited significantly higher stem-driven transpiration (marked ‘A’) than the commercial parent ‘Um-El-Fahem’ (UEF), which showed the lowest recorded rates (marked ‘C’). Mirroring the higher gas exchange activity observed in their stems, the SPC(+) progenies within the segregating population maintained significantly higher transpiration rates (marked ‘A’) compared to the SPC(-) progenies (marked ‘B’). Notably, transpiration rates for all leafless progenies remained consistently above the evaporation baseline measured from empty pots (dotted blue line), confirming that water loss was driven by active physiological regulation within the stems.

**Fig 2.**
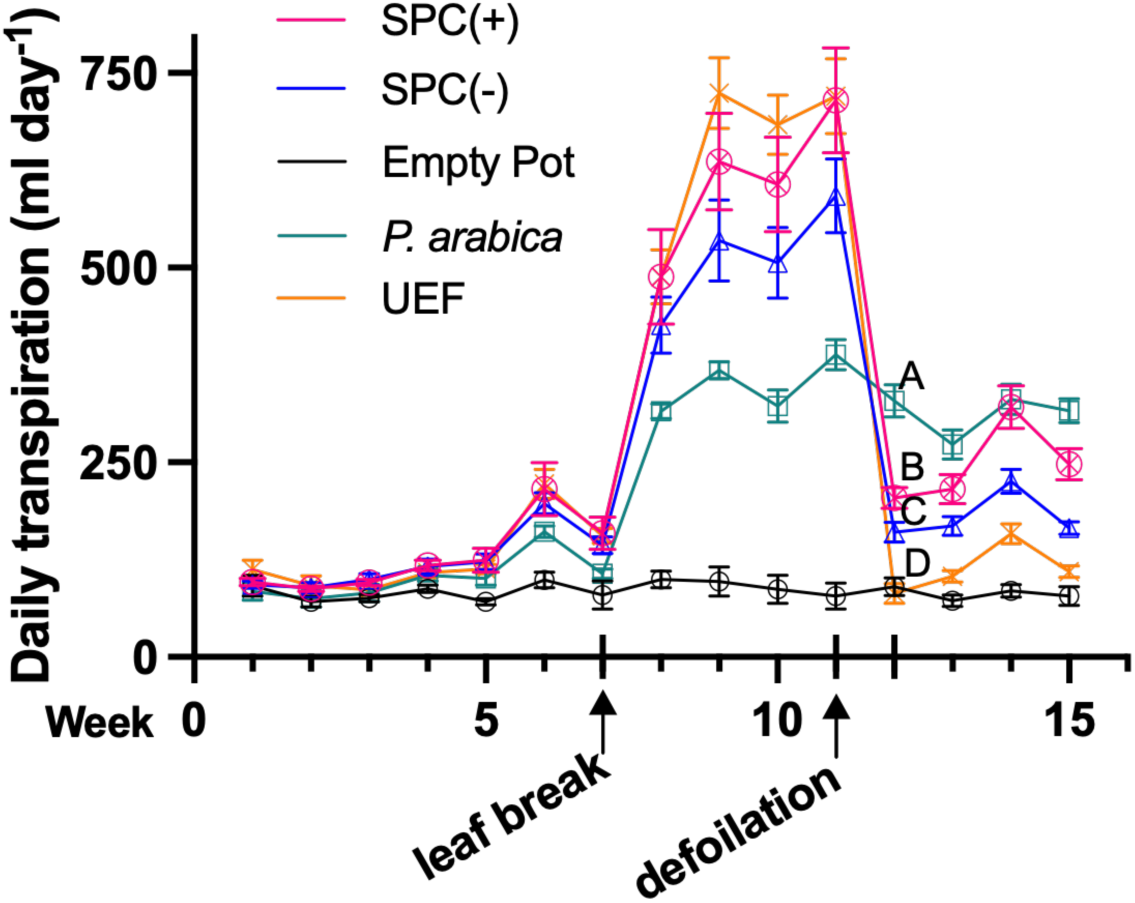
Whole-tree daily transpiration dynamics of ‘Um-El-Fahem’ x *P. arabica* F1 progenies and their parental lines. Data represent daily agronomic transpiration (ml water/day) recorded via the I-CORE lysimetric system starting January 24, 2024 (Week 1). These values were determined by tree daily water consumption via the lysimetric system. Experimental groups include SPC(+) (n = 6 progenies), SPC(-) (n = 6 progenies), and parental lines *P. arabica* (n = 3) and ‘Um-El-Fahem’ (UEF; n = 6). For each progeny, 3-7 biological replicates were measured. Notable events include leaf flush (Week 7) and artificial defoliation (Week 11). The dotted blue line represents the baseline evaporation/conductance measured from empty control pots. Different letters on Week 11 indicate significant differences between experimental groups during the leafless state based on nonparametric Wilcoxon comparisons (*P* < 0.05).

**Fig 3.**
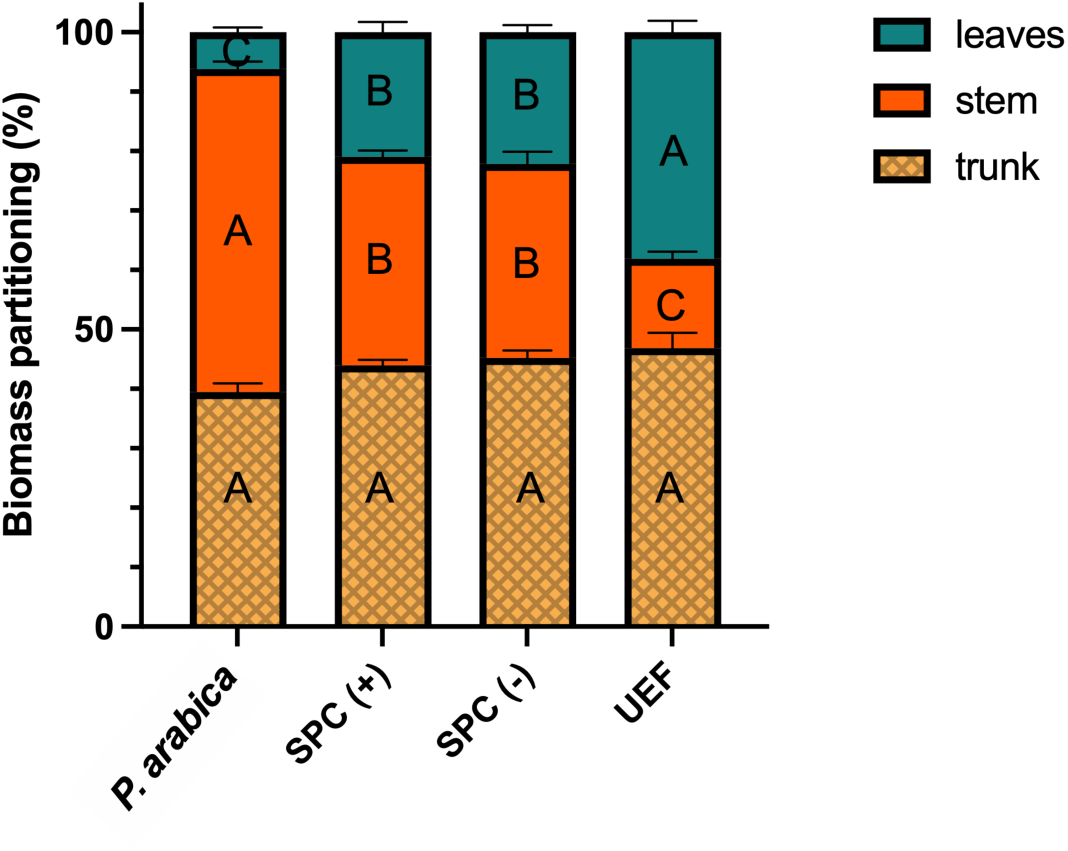
Trunk, stem, and leaf biomass measurements of ‘Um-El-Fahem’ x *P. arabica* F1 progenies and their parental lines. Vegetative dry biomass (g/tree) was measured for twelve selected ‘Um-El-Fahem’ x *P. arabica* F1 progenies (6 SPC(+) and 6 SPC(-)) and their parents. Bars represent the mean dry biomass values, with internal percentages indicating the relative contribution of each tissue (trunk, stem, and leaves). Statistical significance was evaluated using a one-way ANOVA followed by Tukey’s multiple comparisons test to evaluate differences among the four groups independently for each tissue type. Different uppercase letters within a given tissue layer (leaves, stem, or trunk) denote statistically significant differences among the experimental groups (*P* < 0.05). Error bars indicate the standard error of the mean (SEM).

**Fig 4.**
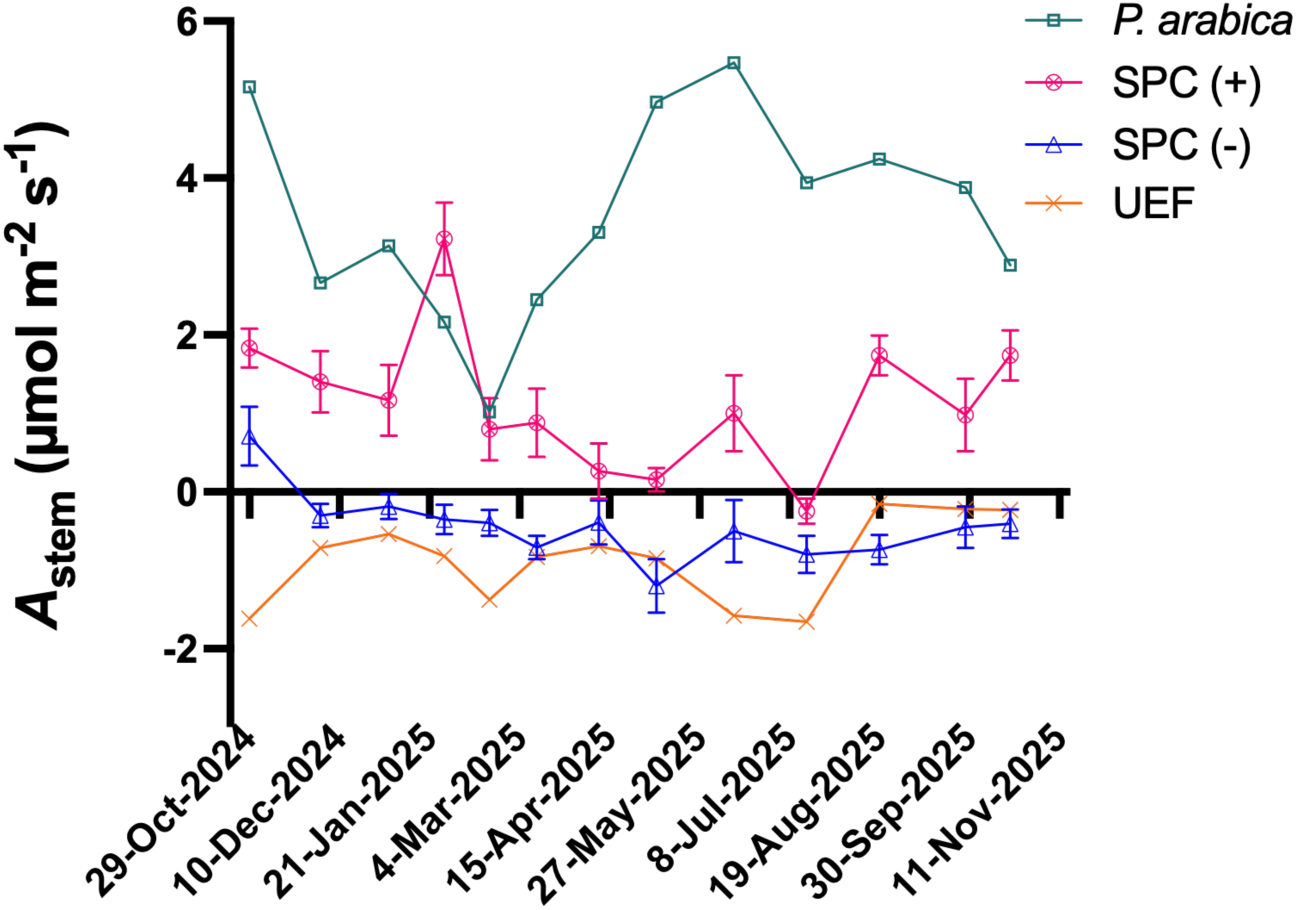
Annual stem CO₂ assimilation dynamics of ‘Um-El-Fahem’ x *P. arabica* F1 progenies and their parental lines. Stem net CO₂ assimilation rates (*A*_stem_) were recorded monthly over a one-year period for twelve selected ‘Um-El-Fahem’ x *P. arabica* F1 progenies (6 SPC(+) and 6 SPC(-)) and their parental lines. Measurements were conducted on four stems per progeny, once a month, over one year. Data points represent the mean *A*_stem_ for each experimental group.

### 3.3 Vegetative Biomass Accumulation and Partitioning

To assess the impact of the SPC trait on growth, vegetative biomass accumulation and partitioning were quantified for all twelve F1 progenies and their parental lines through measurements conducted at two key time points. In mid-April, leaves were harvested to determine fresh and dry weights, while stems and trunks were harvested and weighed at the conclusion of the experiment. Statistical analysis [Nonparametric comparisons for each pair using the Wilcoxon method, *P <* 0.05] revealed that biomass distribution among tissue types differed significantly between groups. Progenies possessing the SPC trait exhibited the highest total biomass (mean = 146 g), which was distributed among the trunk (44%), stems (35%), and leaves (21%). In contrast, progenies lacking the trait (SPC(-)) showed a lower total biomass (mean = 110 g) with a distribution of 46% to the trunk, 32% to the stems, and 22% to the leaves. Among the parental lines, the commercial cultivar ‘Um-El-Fahem’ (UEF) displayed a greater total biomass than the wild species *P. arabica* (mean = 128 g vs. 101 g, respectively), with UEF allocating the largest proportion to the trunk (46%) and leaves (39%) versus smaller values of 15% to the stems, while *P. arabica* dedicated the majority of its biomass to stems (55%), followed by the trunk (39%), and a minimal 6% to leaves.

### 3.4 Annual Stem CO_2_ Assimilation Dynamics in an Orchard Setting

To evaluate the contribution of the stem photosynthetic capability to tree physiology and phenology in their natural environment, we monitor the same progenies that were monitored in a gravimetric lysimetric system, the 6 SPC(+) and 6 SPC(-) *P. arabica* x ‘Um-El-Fahem’ and their parental lines. The implementation of the orchard-based experimental system enabled long-term, multi-year physiological monitoring of mature trees, under natural, uncontrolled field conditions, specifically allowing for the assessment of seasonal phenological traits, including flowering and yield.

To evaluate the year-round carbon assimilation capacity of almond stems, monthly measurements were conducted using a LI-COR 6800 Portable Photosynthesis System on the F1 population and the parental lines grown in the Newe Ya’ar almond orchard. The data revealed distinct net CO_2_ assimilation rates (*A_stem_*) of the tree’s stems between the experimental groups throughout the year. The wild *P. arabica* consistently exhibited the highest *A*_stem_ across all months, while the SPC(+) group showed lower *A*_stem_ than *P. arabica*, but significantly higher values than the SPC(-) and ‘Um-El-Fahem’ (UEF) groups. The SPC(-) and UEF groups displayed nearly identical trends, characterized by negligible or absent *A*_stem_. *A*_stem_ was not uniform throughout the year in any of the groups and exhibited pronounced temporal fluctuations. In *P. arabica*, a marked reduction in *A*_stem_ was observed during dormancy (November-March) yet reached accountable values >2 µmole CO_2_ m^-2^s^-1^ at its lowest values. This was followed by a sharp increase from March-June, after which CO₂ assimilation slowly decreased and stabilized toward the end of the growing season and onset of the next season. In contrast, the SPC(+) group exhibited a unique temporal pattern: a significant increase in *A*_stem_ early in the season (January-February), followed by a decline during the peak leaf-growth period (March-July), and a subsequent gradual increase toward the onset of the next dormancy period (August-November).

### 3.5 Trunk Circumference Analysis

To test whether the Stem Photosynthetic Capability (SPC) trait influenced long-term vegetative growth, trunk circumference was measured at a standardized point, approximately 10 cm above the graft union. This analysis was conducted annually in November over four successive years (2020, 2021, 2023, and 2024) after seasonal tree growth had ceased. The four-year average revealed a significantly larger trunk circumference in trees possessing the SPC trait compared to those lacking it (Fig. 5). Specifically, the SPC(+) group maintained a mean circumference of approximately 44 cm, while the SPC(-) group averaged approximately 33 cm, representing a significant 33.3% increase in trunk secondary growth (*P* < 0.01).

**Fig 5.**
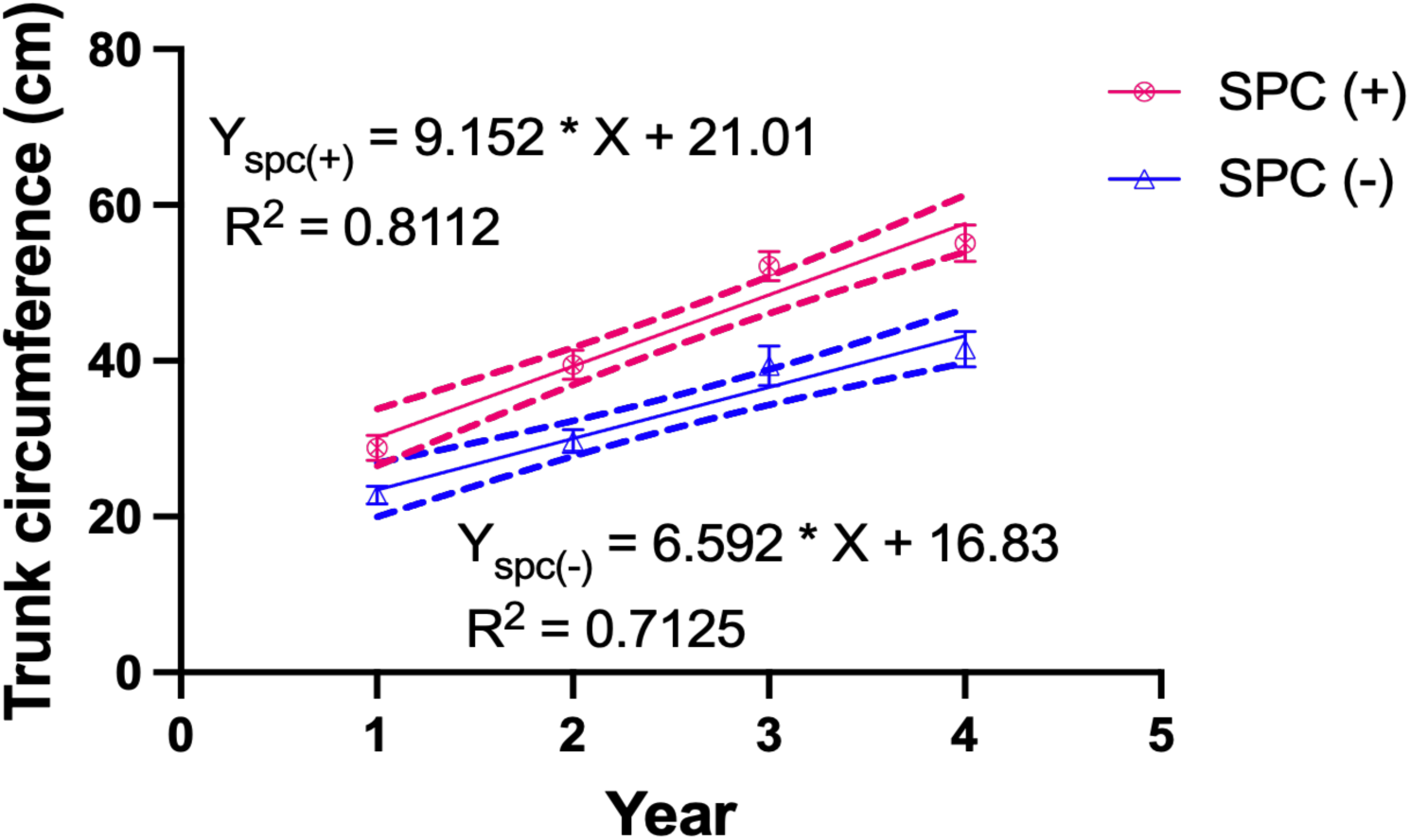
Trunk circumference dynamics in ‘Um-El-Fahem’ x *P. arabica* F1 (2020-2024). Trunk circumference was monitored in twelve selected F1 progenies (n=6 per group). Data represent the annual mean circumference +/-STDEV for the SPC(+) (magenta) and SPC(-) (blue) groups. Solid lines represent linear regressions characterizing the rate of secondary growth, while dotted lines indicate the 95% confidence intervals. Measurements were recorded each November annually following the cessation of vegetative growth. Significant differences in growth trajectories were determined using a Wilcoxon two-sample test with normal approximation (*P* < 0.05).

### 3.6 Blooming Date Analysis

To determine if the Stem Photosynthetic Capability (SPC) affected phenological development, blooming dates were recorded annually from 2021 to 2024. Overall, the population flowered between the beginning of February till early March. To test the influence of SPC on phenology, the date of 10% bloom was recorded for the twelve selected F1 progenies (6 SPC(+) and 6 SPC(-)). Analysis of the four-year average revealed that trees possessing the SPC trait flowered significantly earlier than those lacking it (Fig. 6). Specifically, the SPC(+) group reached the blooming threshold by approximately February 17, whereas the SPC(-) group reached this same developmental stage significantly later, around February 25 (*P* < 0.05).

**Fig 6.**
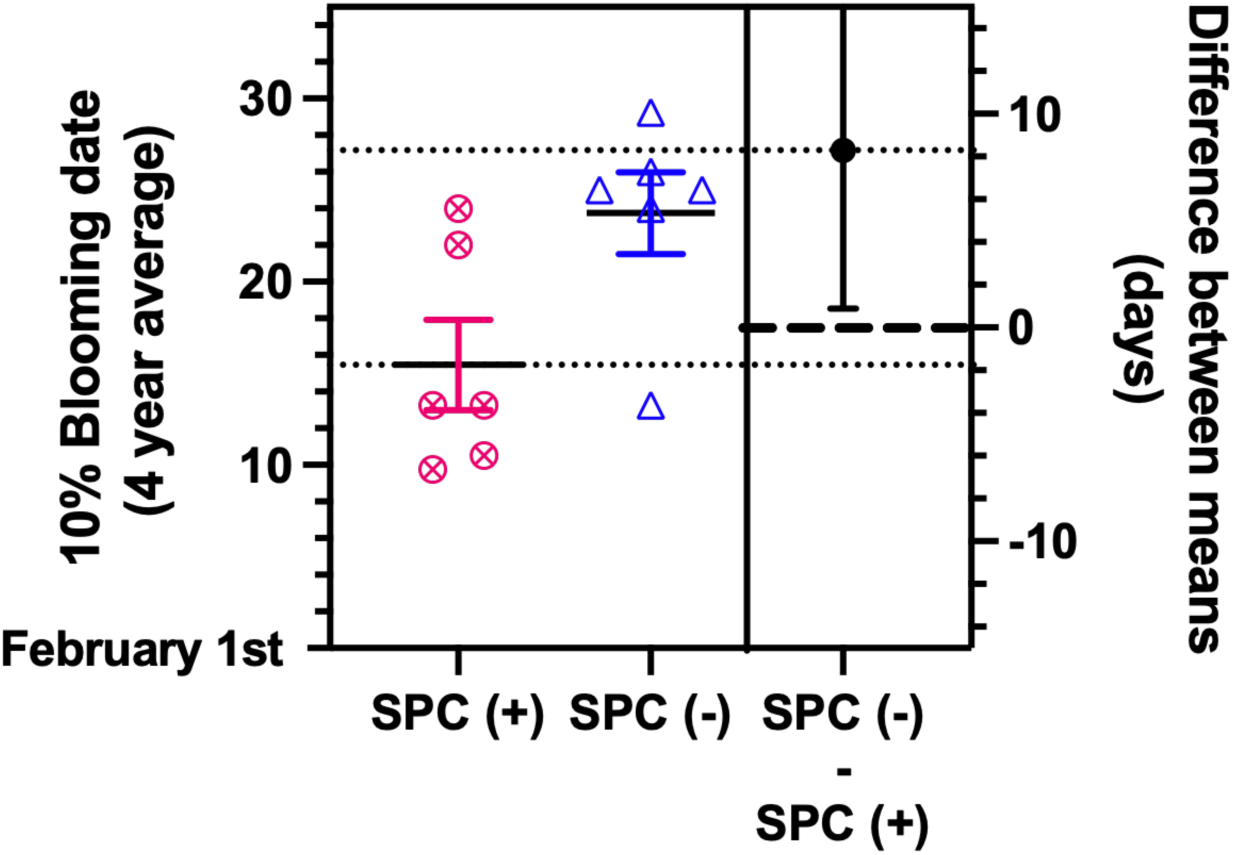
Blooming time of the ‘Um-El-Fahem’ x *P. arabica* F1 progenies from 2020-2024. Blooming time (defined as 10% flowering) was measured in twelve selected ‘Um-El-Fahem’ X *P. arabica* F1 progenies. The analysis was conducted starting February 1^st^ over four successive years (2020-2024). The Y-axis represent the number of days counted from February 1^st^. Data represent the four-year average blooming time +/- STDEV for the two experimental groups (+/- SPC, n = 6 progenies in each group). The mean difference between the SPC(-) (magenta) and SPC(-) (blue) groups is shown on the right floating axes, where the black dot represents the mean difference and the error bar indicates the 95% confidence interval. Statistical significance, determined by a Wilcoxon two-sample test using normal approximation, reveals significant differences between SPC(-) and SPC(+) groups (*P <* 0.05).

### 3.7 Almond Fruit Yield Analysis

To determine if the Stem Photosynthetic Capability (SPC) trait influenced tree productivity, fruit yield was measured in all the ‘Um-El-Fahem’ x *P. arabica* F1 progenies. Ripened fruits were harvested at the end of June. The total average yield, calculated over five successive years (2020-2025), demonstrated a significant positive effect of the of SPC trait on almond productivity (Fig. 7: *P* < 0.01). Specifically, the SPC(+) group achieved a substantially higher mean kernel yield of approximately 2.3 kg per tree, whereas the SPC(-) group averaged significantly lower at approximately 0.5 kg per tree. This robust increase in yield over multiple seasons suggests that the additional carbon contribution from photosynthetic stems directly supports reproductive output in these almond progenies.

**Fig 7.**
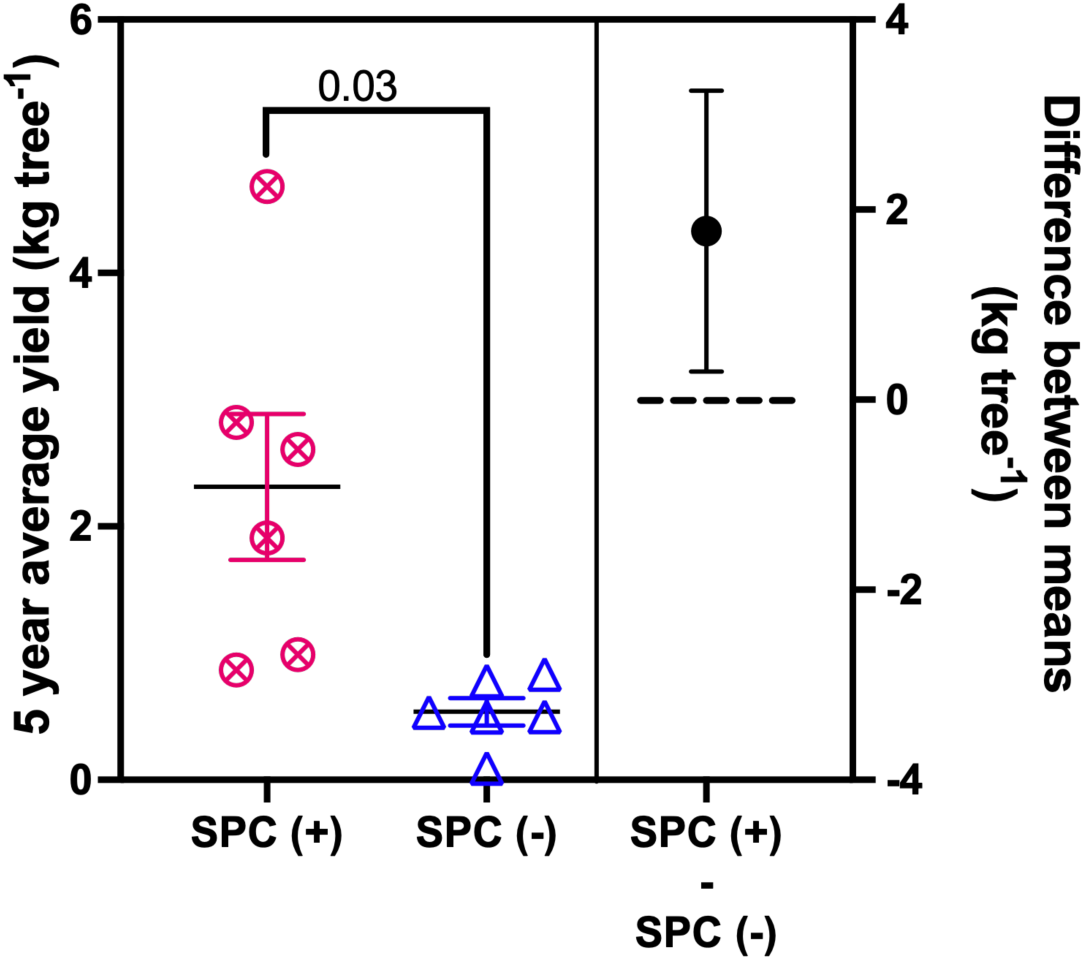
Impact of the SPC trait on almond fruit yield in ‘Um-El-Fahem’ x *P. arabica* F1 progenies. Total kernel weight (kg/tree) was measured for twelve selected F1 progenies (6 SPC(+) and 6 SPC(−)). The average yield of 5 reciprocal years (2020-2025) for the 12 different genotypes, 6 genotypes in each group, SPC(+) (Pink circle), and SPC(-) (Blue triangle), are displayed. The mean difference between the SPC(-) and SPC(-) groups is shown on the right floating axes, where the black dot represents the mean difference and the error bar indicates the 95% confidence interval. Statistical significance, determined by a Wilcoxon two-sample test using normal approximation, reveals significant differences between SPC(-) and SPC(+) groups (*P =* 0.03).

## 4. Discussion

Many adaptive traits found in wild crop relatives have been lost through domestication, yet reintroducing adaptive traits is increasingly critical under current climate change challenges (Dempewolf et al., 2017; Warschefsky et al., 2014). Our study identifies stem photosynthetic capability (SPC) from wild almond *Prunus arabica* as a pivotal trait that is associated with the enhanced whole-tree carbon economy, supports vegetative growth, modulates blooming phenology, and significantly increases yield. By analyzing SPC across an F1 population and its parental lines under varying seasonal and physiological conditions, we demonstrate that this trait is a primary driver of improved physiological performance. These findings highlight SPC as a promising target for improving both the physiological resilience and the agricultural productivity of cultivated almonds.

### 4.1 Integration of Leaf and Stem Carbon Assimilation

To evaluate how SPC contributes to the annual carbon budget, we quantified differences in net CO_2_ assimilation between leaves and stems across genotypes. Our results indicate that leaf carbon assimilation rates (*A*_leaf_) were relatively uniform across all F1 progenies, indicating comparable photosynthetic capability in foliar tissues. In contrast, stem carbon assimilation rates *A*_stem_ varied substantially among groups, particularly during leafless periods, suggesting that variation in whole-tree carbon gain is primarily driven by stem photosynthetic capability rather than leaf performance. Following artificial defoliation, a significant increase in stem carbon assimilation was observed across all genotypes, confirming that stems function as a responsive alternative carbon source. Notably, SPC(+) genotypes exhibited a significantly greater up-regulation of *A*_stem_ compared to SPC(-) genotypes. While the commercial parent ‘Um-El-Fahem’ (UEF) showed only limited stem activity, the wild parent *P. arabica* maintained substantially higher rates, exceeding those of the F1 population. Collectively, these findings are consistent with the conclusion that stem photosynthesis provides a critical, progeny-dependent carbon source that may be vital during periods of reduced leaf activity, such as warm winters or early spring, when respiratory demands remain high but leaf-derived carbon is limited. These differences in stem gas exchange not only impact carbon gain but may also influence whole-tree water dynamics and transpiration.

### 4.2 Physiological Regulation of Whole-Tree Transpiration and Canopy Conductance

Our results (Fig 2) demonstrate that stem photosynthetic capability functions as a dynamic physiological trait, allowing trees to respond flexibly to changing environmental conditions and internal carbon demands. The gravimetric lysimeter platform underscored the functional contribution of SPC to whole-tree physiology, revealing clear differences in water flux among the experimental groups. Because whole-tree transpiration rates are inherently coupled with boundary-layer and tissue conductance across both leaves and stems, these measured water fluxes serve as a direct operational indicator of canopy conductance dynamics.

During the foliated phase, whole-tree transpiration rate and conductance increased across all genotypes following dormancy release and leaf flush. The commercial parent ‘Um-El-Fahem’ (*P. dulcis*) exhibited the highest transpiration rate, consistent with its vigorous growth and high leaf photosynthetic activity, while the wild parent, *P. arabica* displayed lower rates, reflecting its small, thick leaves that are typical of desert-adapted species (Avila-Lovera et al., 2019; Trainin et al., 2022). Among the ‘Um-El-Fahem’ x *P. arabica* F1 progenies, SPC(+) progenies showed significantly higher conductance than SPC(-) progenies, reflecting the supplemental contribution of active stem photosynthetic tissues (i.e., SPC trait) to the overall gas exchange of the tree canopy.

Following artificial defoliation, marked physiological differences emerged: SPC(+) progenies maintained high canopy conductance, tracked directly via elevated stem transpiration rates, comparable to *P. arabica*, whereas SPC(-) progenies and the commercial parent ‘Um-El-Fahem’ exhibited significantly lower levels. This demonstrates that, in the absence of leaves, whole-tree transpiration is driven almost exclusively by active stem tissues, as consistently observed in *P. arabica* and the SPC(+) group, which exhibited significantly higher rates compared to SPC(-) and ‘Um-El-Fahem’. Because transpiration and photosynthesis are tightly coupled via stomatal regulation, the enhanced stem-driven canopy conductance can be used as a physiological proxy for increased stem carbon assimilation (Avila-Lovera et al., 2017; Nilsen and Bao, 1990). We therefore quantified the contribution of the SPC trait to vegetative biomass accumulation and its partitioning within the tree’s aerial architecture (i.e., trunk, stems, and leaves).

### 4.3 Biomass Partitioning and Carbon Allocation Strategies

The parental lines utilized in this study displayed divergent strategies for biomass allocation across their primary aerial tissues, namely the trunk, stems, and leaves. *P. arabica* directs the majority of its vegetative biomass toward green stems (54.3%), a structural investment that reflects its high stem photosynthetic capability. In contrast, ‘Um-El-Fahem’ (UEF) primarily allocates resources to the trunk (48.6%) and leaves (38%), with minimal investment (15%) in stem tissue. These distinct patterns highlight species-specific mechanisms for maximizing carbon gain, with each parent prioritizing the development of the tissues that contribute most significantly to their respective photosynthetic strategies.

Among the F1 progenies, biomass allocation appears to represent an intermediate strategy, combining features from both parental lines. Both SPC(+) and SPC(-) progenies allocated a substantial and comparable portion of their biomass to the trunk (∼44–46%), while maintaining intermediate levels of investment in stems and leaves. Although SPC(+) individuals exhibited higher total biomass than SPC(-) progenies (mean = 146 g vs. 110 g, respectively), these differences were not statistically significant. This pattern likely results from their shared origin as a segregating F1 population, in which each progeny inherits a mixture of traits from both parents, moderating the effect of stem photosynthetic capability on overall biomass distribution. Nevertheless, the results suggest that the SPC trait in the F1 progenies may enhance whole-tree carbon gain while allowing the trees to maintain a balanced partitioning strategy across the various aerial tissues (i.e., trunk, stems, and leaves). This integrated approach reflects a compromise between the specialized carbon acquisition strategies of *P. arabica* and ‘Um-El-Fahem’.

### 4.4 Annual Stem CO_2_ Assimilation Dynamics in an Orchard Setting

Monitoring stem CO_2_ assimilation within the orchard environment revealed distinct differences in how the SPC(+) and SPC(-) progeny groups utilized their stems across seasonal cycles throughout the year. The wild parent *P. arabica* consistently maintained higher stem CO_2_ assimilation capacity compared to both the F1 progenies and the commercial parent ‘Um-El-Fahem,’ even during periods when leaves were present. Among the F1 population, SPC(+) group exhibited intermediate yet significant stem CO_2_ assimilation activity, which was markedly higher than SPC(-) group and the commercial parent, both of which displayed minimal to negligible stem-based carbon fixation.

Notably, stem assimilation was not constant throughout the year and exhibited pronounced temporal fluctuations in all experimental groups. In SPC(+) progenies, stem net CO_2_ assimilation rates increased substantially during leafless periods and reached a pronounced peak in February, coinciding with the flowering period. This seasonal pattern is consistent with evidence that green stems can sustain carbon assimilation when foliage is absent and make an important contribution to whole-plant carbon balance and carbohydrate reserves (Avila-Lovera et al., 2019; Natale et al., 2023). This temporal synchronization is consistent with the idea that photosynthetic stems may serve as a critical supplementary carbon source during energetically demanding developmental stages (Avila-Lovera et al., 2024; Natale et al., 2023; Simkin et al., 2020). Following leaf emergence, stem net CO_2_ assimilation rates declined, indicating a dynamic redistribution of photosynthetic activity between leaves and stems as canopy development progresses and tissue availability changes. This observation is consistent with studies showing seasonal shifts in the relative contribution of stems and leaves to carbon gain (Aschan and Pfanz, 2003; Tinoco-Ojanguren, 2008). Overall, these results demonstrate that SPC may provide vital temporal flexibility in carbon supply, supporting energy-intensive processes when foliar photosynthesis is limited, a physiological advantage likely inherited from *P. arabica* and consistent with evidence that green stems can sustain carbon gain and non-structural carbohydrate reserves under stressful conditions (Avila-Lovera et al., 2017; Natale et al., 2023; Simkin et al., 2020).

### 4.5 Effects of SPC on Long-Term Vegetative Growth

One of the most important indicators of the functional relevance of SPC is its effect on tree growth. Additional CO₂ assimilation through photosynthetic stems can enhance whole-plant carbon gain, supporting biomass accumulation and survival, particularly under stress (Avila-Lovera et al., 2024; Avila-Lovera et al., 2019; Pfanz et al., 2002). To assess these long-term growth differences, trunk circumference was measured annually at a standardized point in the almond orchard over four consecutive years, providing a comprehensive multi-year perspective on tree development. Unlike the biomass measurements in the lysimeter-based experiment, which reflected a shorter duration and showed no significant differences, the multi-year, orchard analysis revealed a clear and significant increase in trunk circumference in SPC(+) progenies compared to SPC(-) individuals. Specifically, the SPC(+) group achieved a significant 33.3% increase in trunk secondary growth. This divergence is likely attributable to the presence of stem tissue capable of assimilating CO_2_ during periods when foliage is absent (such as winter or early spring in deciduous fruit trees) or during photosynthetic-limited periods, with green stems known to maintain carbon assimilation and non-structural carbohydrate supply under such conditions (Avila-Lovera et al., 2017; Natale et al., 2023). By supplying additional carbon during periods when leaf photosynthesis is limited, SPC(+) progenies can allocate more energy to trunk thickening and overall vegetative growth, resources that remain inaccessible to SPC(-) progenies. These findings demonstrate that stem photosynthetic capability contributes not only to short-term carbon gain but also to cumulative biomass accumulation over multiple years, consistent with studies showing that green stems enhance plant carbon balance and growth (Avila-Lovera et al., 2019; Pfanz et al., 2002; Ye et al., 2025).

### 4.6 Effects of SPC on Phenological Progression and Blooming Time

Analysis of the blooming time index over four consecutive years confirmed that the SPC trait consistently advances reproductive phenology. SPC(+) trees reached the 10% bloom stage sooner than their SPC(-) counterparts, suggesting that the supplemental carbon provided by active stem tissues may facilitate an earlier initiation of flowering. While the SPC(-) group reached the blooming threshold 23 days after February 1^st^, the SPC(+) group reached it in just 15 days. This phenological shift is consistent with evidence that photosynthetically active stems can sustain carbon assimilation and carbohydrate reserves when foliage is absent or limited, thereby supporting metabolically demanding developmental transitions (Avila-Lovera et al., 2024; Natale et al., 2023). This extra carbon can modify the balance of soluble sugars within the tree, which may influence complex biological processes such as dormancy release and progression to flowering (Chmielewski and Gotz, 2022; Kaufmann and Blanke, 2017; Sperling et al., 2019; Tixier et al., 2019). By increasing sugar availability, SPC(+) progenies can progress more efficiently to the flowering stage, reaching the 10% flowering threshold without heavily drawing on stored reserves. In practical terms, this early progression into flowering could be advantageous under future climate scenarios, where warmer winters are expected to reduce winter chill accumulation and jeopardize timely bloom in many deciduous fruit and nut trees (Luedeling et al., 2011; Sperling et al., 2019). In such contexts, SPC(+) trees may better maintain timely flowering and high yield potential because stem photosynthesis can sustain carbohydrate reserves and hydraulic function when foliar photosynthesis is constrained (Natale et al., 2023). This progression highlights the functional role of SPC not only in growth but also in coordinating reproductive phenology.

### 4.7 Cumulative effect on yield

The yield data represent the total kernel weight produced by all trees over five years (2020-2025), and the multi-year average clearly reflects differences between the experimental groups. SPC(+) group consistently achieved a markedly higher kernel yield than SPC(-) group, a difference that remained statistically significant throughout every year of the study. Specifically, the SPC(+) group maintained a substantially higher mean kernel yield of approximately 2.3 kg per tree, whereas the SPC(-) group averaged only 0.5 kg per tree. This persistent 4.6-fold advantage indicates that stem photosynthetic capacity provides additional carbon that can be invested in reproductive structures, consistent with studies showing that stem and other non-foliar photosynthesis can contribute meaningfully to plant growth and yield (Avila-Lovera et al., 2024; Natale et al., 2023; Simkin et al., 2020). The presence of additional photosynthetic tissue in SPC(+) trees allows for CO_2_ assimilation even during the dormant season, providing a supplemental carbon source that is unavailable to genotypes lacking the trait. These winter-assimilated carbon reserves can then be mobilized during the growing season to support essential processes such as flowering, fruit set, and kernel development, as non-structural carbohydrates and sugar signaling are known to play central roles in reproductive development and yield formation in fruit trees (Rossouw et al., 2024; Zwieniecki et al., 2022). Furthermore, photosynthetically active stems may play a vital role during the early stages of fruit development, when leaf photosynthetic capacity is still limited and metabolic carbon demand is at its peak (Avila-Lovera et al., 2019; Ye et al., 2025). Consequently, stem photosynthesis contributes not only to vegetative growth and carbon balance but also to a stable, long-term increase in reproductive output, highlighting its significant ecological and agronomic value in almond breeding.

## Acknowledgements

We thank Prof. Menachem Moshelion (Faculty of Agriculture, The Hebrew University of Jerusalem, Rehovot, Israel) for supporting and supervising the lysimetric experimental and analytical work.

## CRediT Authorship Contribution Statement

**Dan Zeira:** Investigation, Formal analysis, Project administration, Visualization, Writing – original draft. **Ori Eisenbach:** Investigation, Formal analysis. **Rotem Harel-Beja:** Investigation, Resources. **Taly Trainin:** Investigation, Resources. **Kamel Hatib:** Resources. **Lotem Terner:** Resources. **Mohamed Abd-Elhadi:** Resources. **Hillel Brukental:** Investigation, Formal analysis. **Or Shapira:** Conceptualization, Writing - review & editing. **Yotam Zait:** Conceptualization, Writing - review & editing. **Doron Holland:** Conceptualization, Writing – review & editing. **Tamar Azoulay-Shemer:** Conceptualization, Methodology, Supervision, Project administration, Funding acquisition, Visualization, Writing – review & editing.

## Conflict of interest

The authors declare no conflict of interest.

## Funding

This work was supported by the Ministry of Innovation, Science & Technology, Israel (#1001577995), by the State of Israel Ministry of Agriculture Rural Development & Chief Scientist Office (#20-01-0271), and by the “Leona M. and Harry B. Helmsley Charitable Trust”.

## Data availability

All primary data to support the findings of this study are openly available in the Zenodo Repository, https://shorturl.at/kQHuc

## Notes

https://shorturl.at/kQHuc

### Competing Interest Statement

The authors have declared no competing interest.

